# Improving polygenic prediction from summary data by learning patterns of effect sharing across multiple phenotypes

**DOI:** 10.1101/2024.05.06.592745

**Authors:** Deborah Kunkel, Peter Sørensen, Vijay Shankar, Fabio Morgante

## Abstract

Polygenic prediction of complex trait phenotypes has become important in human genetics, especially in the context of precision medicine. Recently, Morgante *et al*. introduced *mr.mash*, a flexible and computationally efficient method that models multiple phenotypes jointly and leverages sharing of effects across such phenotypes to improve prediction accuracy. However, a drawback of *mr.mash* is that it requires individual-level data, which are often not publicly available. In this work, we introduce *mr.mash-rss*, an extension of the *mr.mash* model that requires only summary statistics from Genome-Wide Association Studies (GWAS) and linkage disequilibrium (LD) estimates from a reference panel. By using summary data, we achieve the twin goal of increasing the applicability of the *mr.mash* model to data sets that are not publicly available and making it scalable to biobank-size data. Through simulations, we show that *mr.mash-rss* is competitive with, and often outperforms, current state-of-the-art methods for single- and multi-phenotype polygenic prediction in a variety of scenarios that differ in the pattern of effect sharing across phenotypes, the number of phenotypes, the number of causal variants, and the genomic heritability. We also present a real data analysis of 16 blood cell phenotypes in UK Biobank, showing that *mr.mash-rss* achieves higher prediction accuracy than competing methods for the majority of traits, especially when the data has smaller sample size.

**Author summary:** Polygenic prediction refers to the use of an individual’s genetic information (*i.e*., genotypes) to predict traits (*i.e*., phenotypes), which are often of medical relevance. It is known that some phenotypes are related and are affected by the same genotypes. When this is the case, it is possible to improve the accuracy of predictions by using methods that model multiple phenotypes jointly and account for shared effects. *mr.mash* is a recently developed multi-phenotype method that can learn which effects are shared and has been shown to improve prediction. However, *mr.mash* requires large data sets of genetic and phenotypic information collected at the individual level. Such data are often unavailable due to privacy concerns, or are difficult to work with due to the computational resources needed to analyze data of this size. Our work extends *mr.mash* to require only summary statistics from Genome-Wide Association Studies instead of individual-level data, which are usually publicly available. In addition, the computations using summary statistics do not depend on sample size, making the newly developed *mr.mash-rss* scalable to extremely large data sets. Using simulations and real data analysis, we show that our method is competitive with other methods for polygenic prediction.

## Introduction

Predicting complex trait phenotypes from genotypes is a central task of a few branches of quantitative genetics. In agricultural breeding, there is interest in predicting breeding values (EBV) to select the best individuals for reproduction and achieve an increase in performance over generations [1]. In human genetics, predicting medically relevant phenotypes such as disease risk via the so-called polygenic scores (PGS) is important to stratify the population and identify individuals with greater genetic risk [2]. Finally, with the advent of transcriptome-wide association studies (TWAS), predicting gene expression as an intermediate step has become of interest [3]. In all these applications, accurate predictions are important. The response to artificial selection is directly proportional to the accuracy of EBVs [4]. Precise identification of individuals at risk for a particular disease requires accurate PGS [2]. The power to discover gene-phenotype associations in TWAS depends on the accuracy of gene expression prediction [5].

Technically, phenotypic prediction is achieved by modeling the phenotype of interest as a multiple regression on genotypes at a set of genetic variants [6]. Both frequentist and Bayesian approaches to multiple regression have been developed for and/or applied to this task, with accuracy spanning from very low to high depending on the genetic architecture of the trait analyzed [7–12]. Multiple phenotypes may be genetically correlated due to pleiotropy (*i.e*., the sharing of causal variants across traits). In that case, modeling these phenotypes jointly via multivariate multiple regression methods can improve effect sizes estimates by leveraging effect sharing and, thus, increase prediction accuracy [13–17].

Recently, Morgante *et al*. (2023) introduced the “Multiple Regression with Multivariate Adaptive Shrinkage” or “*mr.mash*” [18]. *mr.mash* is a Bayesian approach to multivariate multiple regression that is able learn complex patterns of effect sharing across phenotypes directly from the data. This is achieved through the use of flexible priors on the effect sizes across phenotypes and an empirical Bayes framework to adapt these priors to the data. Computational effiency is achieved by using Variational Inference (VI) as opposed to the more expensive Markov Chain Monte Carlo (MCMC) methods. Using multi-tissue gene expression prediction from cis-genotypes as an example, the authors showed that *mr.mash* is competitive in terms of both prediction accuracy and speed [18]. However, while powerful, *mr.mash* has some limitations. First, *mr.mash* requires individual-level data, *i.e*., genotypes and phenotypes for each individual and, mainly for privacy reasons, these data are rarely publicly available [19]. Second, *mr.mash* does not scale well to datasets with very large sample size, such as modern biobanks. These weaknesses limit the use of *mr.mash* for PGS prediction in human genetics.

In this work, we overcome both these limitations by introducing “*mr.mash* Regression with Summary Statistics” or “*mr.mash-rss*”, an extension of *mr.mash* that only requires summary-level data. These are effect sizes and their standard error (or Z-scores) from univariate Genome-Wide Association Study (GWAS) and Linkage Disequilibrium estimates from reference panels, which are usually publicly available [19]. We test *mr.mash-rss* in the task of PGS prediction for multiple phenotypes jointly via simulations in several scenarios and show that it is competitive in terms of prediction accuracy with currently available methods. We then confirm these results in the analysis of real data for 16 blood cell traits in the UK Biobank [20, 21].

### Description of the method

The multivariate multiple regression is used to model the effects of several predictor variables ***X*** on multiple responses ***Y*** jointly:

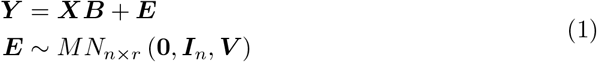

where ***Y*** ∈ ℝ^*n×r*^ is the response matrix for *r* responses (phenotypes in our case) in *n* individuals, ***X*** ∈ ℝ^*n×p*^ is the predictor matrix for *p* predictors (genetic variants in our case) in *n* individuals, ***B*** ∈ ℝ^*p×r*^ is the matrix of effects for *p* predictors and *r* responses, and ***E*** ∈ ℝ^*n×r*^, is the matrix of residuals for *r* responses for *n* individuals. The residuals follow a Matrix Normal distribution with mean **0** ∈ ℝ^*n×r*^, covariance across individuals ***I***_*n*_ ∈ ℝ^*n×n*^ (an identity matrix), and covariance across responses ***V*** ∈ ℝ^*r×r*^.

*mr.mash* adopts a Bayesian approach by imposing a prior on the effects:

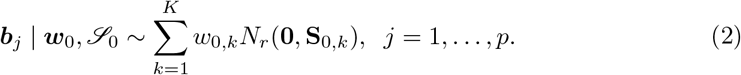

where ***b***_*j*_ is an *r*-vector that captures the effects of predictor *j*, and 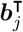 is the *j*^*th*^ row of ***B***. Thus, the effects are assumed to be identically distributed as a mixture of *r*-variate normals with *K* components. The prior is determined by ***w***_0_ := (*w*_0,1_, …, *w*_0,*K*_), the set of non-negative mixture weights, and ℐ_0_ := {***S***_0,1_, …, ***S***_0,*K*_ }, the set of *r × r* covariance matrices across responses. The elements of ℐ_0_ are prespecified and are intended to capture plausible patterns of effect sharing across responses [18].

To make the model fit computationally efficient for large datasets, *mr.mash* approximates *p*(***B*** | ***X, Y***, ***V***, ***w***_0_, ℐ_0_), the true posterior distribution of the regression coefficients, through variational inference, which uses optimization techniques to find the best approximation within a chosen family of distributions [22]. The optimal approximation is determined by maximizing the evidence lower bound (ELBO), a lower bound on the model’s marginal likelihood. In addition, *mr.mash* also estimates ***w***_0_ (and ***V***) from the data by maximizing the ELBO, thereby adapting the prior to the data. This whole procedure has been termed variational empirical Bayes [23].

### Extension of *mr.mash* to summary statistics

Following the approach of [24], we express the updates in *mr.mash* in terms of sufficient statistics. The likelihood for the *mr.mash* model is

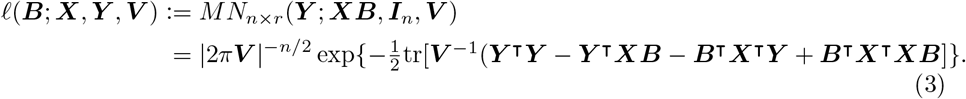

We can see that ***X***^⊺^***X, X***^⊺^***Y***, and ***Y*** ^⊺^***Y*** are sufficient statistics for the likelihood. Thus, the *mr.mash* model can be fitted using expressions based only on these sufficient statistics (see S1 Text for detailed derivations) and obtain the same results as using individual-level data ***X*** and ***Y***.

The sufficient statistics can be recovered from effect sizes and their standard error (or Z-scores) from GWAS and LD estimates, following steps provided in [24] and S1 Text. We call *mr.mash* with summary statistics *mr.mash-rss*. However, it should be noted that while ***X***^⊺^***Y*** can be recovered exactly, ***X***^⊺^***X*** is only approximated when LD estimates come from reference panels, rather than from the data that generated GWAS summary statistics [24]. Thus, using summary data can be seen as fitting the *mr.mash* model using an approximation to the likelihood in 3 [24]. The quality of the approximation depends on how closely the LD reference panel matches the GWAS summary statistics. Quality control should therefore be performed on summary statistics and LD before model fitting [24, 25]. In addition, ***Y*** ^⊺^***Y*** may not be available. However, this quantity is not strictly necessary, unless ***V*** is estimated within the *mr.mash-rss* algorithm [24]. While *mr.mash* has a way to deal with missing values in ***Y***, *mr.mash-rss* assumes the summary statistics be computed using the same individuals for each response (*i.e*., there are no missing values in ***Y***).

### Software availability

The methods introduced in this paper are implemented in the R package [26] `mr.mash.alpha` which is available for download at https://github.com/stephenslab/mr.mash.alpha.

## Verification and comparison

### Simulations using UK Biobank genotypes

We devised a simulation study where the goal was to compare *mr.mash-rss* and other competing methods at computing PGS for multiple phenotypes from summary data. We used real genotypes from the UK Biobank array data for *n* =105,000 nominally unrelated White British individuals that were randomly sampled. After applying a series of filters (see S1 Text for details), the data included *p* =595,071 genetic variants.

We simulated *r* = 5 phenotypes according to three scenarios that differed in the structure of the effect sharing across phenotypes. Causal variants (5,000 for all scenarios) were randomly sampled from all genetic variants.

1. “Equal Effects”, where each causal variant affects all the phenotypes and has the same effect across phenotypes. The per-phenotype proportion of variance explained by the causal variants or genomic heritability 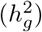 is equal to 0.5.
2. “Mostly Null”, where the causal variants affect only the first phenotype with 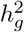 equal to 0.5, while the remaining phenotypes are affected only by an non-genetic component (*i.e*., 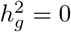)
3. “Shared Effects in Subgroups”, where the effect of each causal variant is drawn such that it is equally likely to be shared (but not be equal) in phenotypes 1 through 3 or to be shared (but not be equal) in phenotypes 4 and 5. The per-phenotype 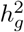 is 0.3 in phenotypes 1-3 and 0.5 is phenotypes 4 and 5. These three scenarios were similar to those used in [18], but some parameters (e.g., number of causal variants) were modified to reflect more closely the genetic architecture of complex traits, rather than gene expression. We also simulated a few scenarios based on the Equal Effects scenario (*i.e*., equal effects of the causal variants across phenotypes) to assess the effect of genomic heritability, polygenicity (*i.e*., number of causal variants), and number of phenotypes modeled on the performance of the methods:
4. “Low 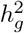”, where the per-phenotype 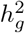 is 0.2.
5. “High Polygenicity”, where the number of causal variants is 50,000.
6. ”More Phenotypes”, where the number of simulated phenotypes is 10.

For each of the scenarios above, we simulated 20 replicates. Per-phenotype prediction accuracy was computed as the *R*^2^ from the linear regression of the true phenotypes on the predicted phenotypes for the test set individuals, which consisted of 5,000 randomly sampled individuals from the total of 105,000. This metric has the attractive property that its upper bound is 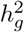 [27].

### Methods compared

We compared *mr.mash-rss* to a few competing methods that satisfied the following requirements: (1) can be fitted with only summary data; (2) do not require a validation data set to tune model parameters; (3) for multivariate methods, are able to model at least 5 phenotypes *jointly*. This resulted in the choice of the following methods:

1. *LDpred2-auto*. This is a univariate Bayesian method that imposes a two-component mixture prior on the regression coefficients, consisting of a point-mass at 0 and a zero-centered normal distribution [10]. This method is labelled “LDpred2” in the results.
2. *SBayesR*. This is a univariate Bayesian method that imposes a four-component mixture prior on the regression coefficients, consisting of a point-mass at 0 and three zero-centered normal distributions, each with a different variance [9]. This method is labelled “SBayesR” in the results.
3. *SmvBayesC*. This is a multivariate Bayesian method that imposes a two-component mixture prior on the regression coefficients across phenotypes, consisting of a point-mass at 0 and a zero-centered multivariate normal distribution [28, 29]. This method allows for each genetic variant to affect any combination of phenotypes. This method is labelled “SmvBayesC” in the results. We also tested a “restrictive” version that allows for each genetic variant to affect all or none of the phenotypes only [14, 28]. This method is labelled “SmvBayesC-rest” in the results.

Each method was fitted for each chromosome separately using summary statistics calculated using only the training set individuals. The summary statistics (*i.e*., effect sizes and standard errors) were computed from univariate simple linear regression of each phenotype on each genetic variant, one at a time. Each phenotype was quantile normalized before the analysis. LD between each pair of variants was computed using 146,288 nominally unrelated White British individuals that did not overlap with the 105,000 individuals used for the rest of the analyses. Correlations between variants that were more than 3 cM apart were set to 0 to create a sparse LD matrix [10]. We fitted *mr.mash-rss* including both “canonical” and “data-driven” covariance matrices (see S1 Text and [18] for details).

## Results

In the Equal Effects scenario (Fig 1A), the three multivariate methods performed better than the two univariate methods. This is expected because the univariate methods assume independence among genetic effect across phenotypes and are unable to learn the pattern of equal genetic effects. Among the multivariate methods, *mr.mash-rss* produced higher accuracy than *SmvBayesC*. The “restrictive” version of *SmvBayesC* performed as well as the unrestricted one because this scenario meets one of the effect combinations allowed by this less flexible method.

**Fig 1.**
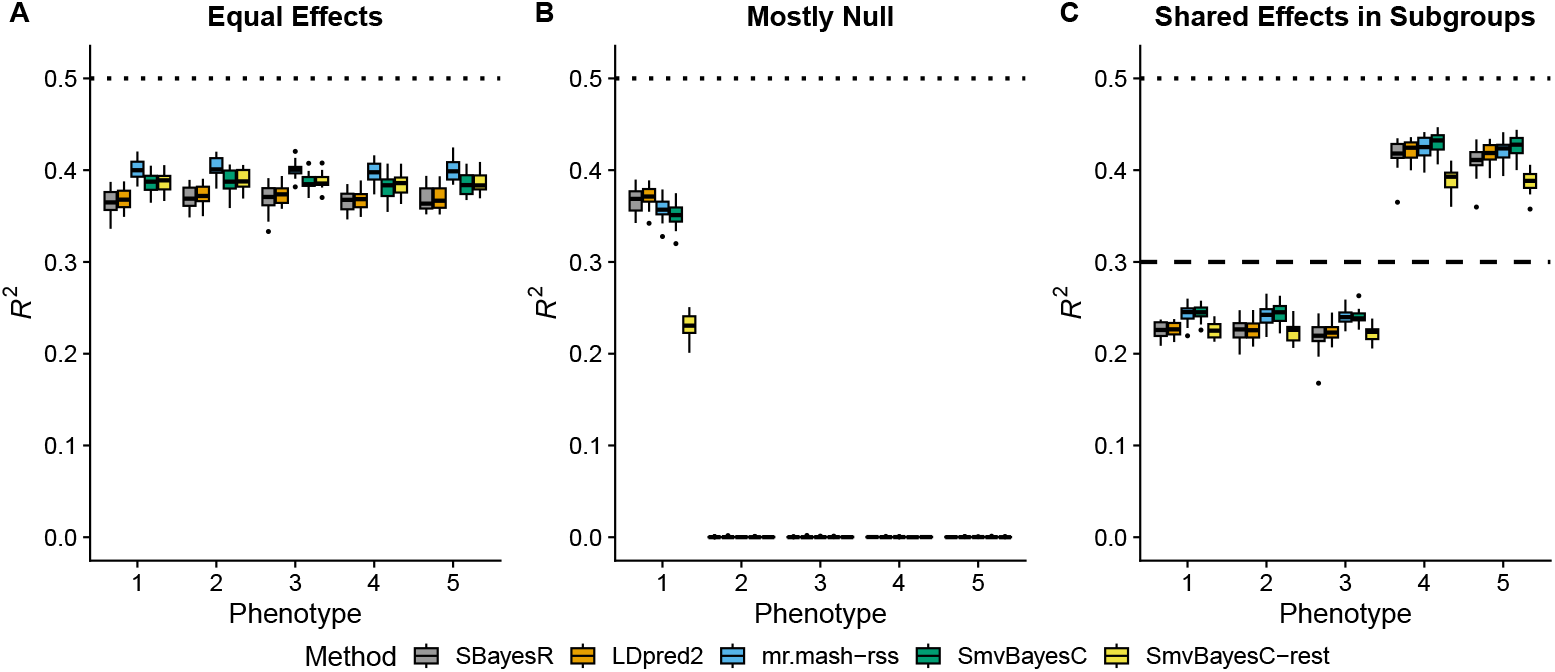
Prediction accuracy in simulations with different patterns of effect sharing across phenotypes. Each panel summarizes the accuracy of the test set predictions in 20 simulations. The thick, black line in each box gives the median *R*^2^. The dotted and dashed lines give the maximum accuracy achievable, *i.e*., the simulated 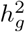.

In the Mostly Null scenario (Fig 1B), the genetic effects are present only in the first phenotype. Thus, joint modeling of all the phenotypes is not expected to produce any increase in accuracy compared to a phenotype-by-phenotype analysis. In phenotype 1, while *SBayesR* and *LDpred2-auto* were the most accurate methods, *mr.mash-rss* only had slightly lower mean *R*^2^. As for *SmvBayesC*, the full version performed only slightly worse than *mr.mash-rss*; however, the “restrictive” version performed much worse. This observation was expected, given that the prior of *SmvBayesC* “restrictive” only allows for the effects to be present in all or none of the phenotypes.

In the Shared Effects in Subgroups scenario (Fig 1C), *SBayesR, LDpred2-auto, SmvBayesC*, and *mr.mash-rss* performed very similarly in phenotypes 4 and 5, with *SmvBayesC* having slightly higher accuracy than the other methods. On the other hand, *SmvBayesC* and *mr.mash-rss* outperformed the univariate methods in phenotypes 1-3. This can be explained by the slightly higher sharing of effects across phenotypes, the larger number of phenotypes with shared effects, and the lower 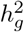, which make the advantage of a multivariate analysis more clear than in phenotypes 4 and 5. The prior of *SmvBayesC* “restrictive” is not well suited for this scenario, which results in this method being the worst across phenotypes.

In the Low 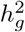 scenario (Fig 2A), the three multivariate methods performed better than the two univariate methods. In addition, the relative improvement provided by the multivariate methods is larger than in the Equal Effects scenario with 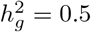 (Fig 1A). With smaller signal-to-noise ratio, it is harder to estimate effects accurately. Multivariate methods can borrow information across phenotypes and improve accuracy. *mr.mash-rss* is the best performing method in this scenario.

**Fig 2.**
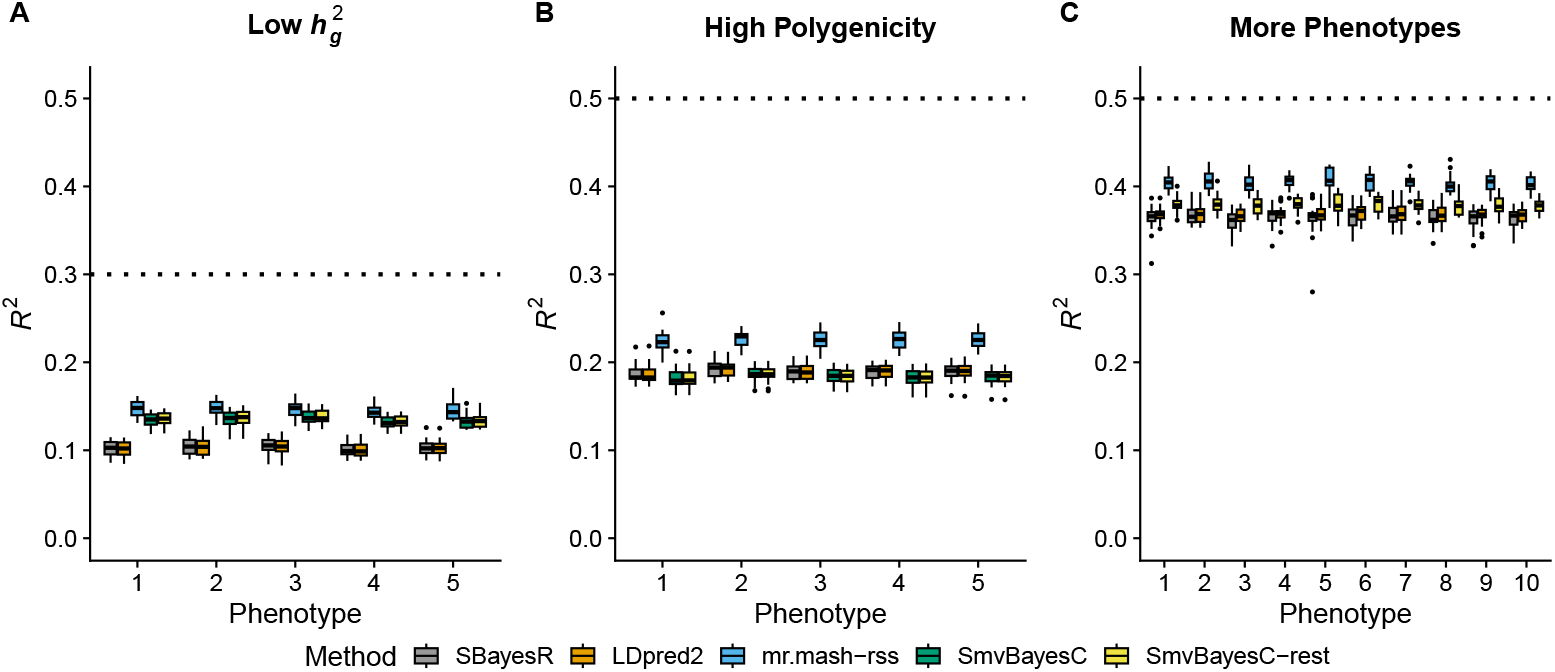
Prediction accuracy in simulations with different genetic architecture. Each panel summarizes the accuracy of the test set predictions in 20 simulations. The thick, black line in each box gives the median *R*^2^. The dotted lines give the maximum accuracy achievable, *i.e*., the simulated 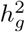.

In the High Polygenicity scenario (Fig 2B), prediction accuracy achieved by all the methods is much lower than in the Equal Effects scenario with 5,000 causal variants (Fig 1A). This is expected since each causal variant explains a much smaller proportion of phenotypic variance and, consequently, the effects are harder to estimate accurately. However, *mr.mash-rss* substantially outperformed both univariate and multivariate competing methods. *SmvBayesC* could not adapt well to this scenario, providing accuracies that are similar to or even lower than *SBayesR* and *LDpred2-auto*.

In the More Phenotypes scenario (Fig 2C), the results are very similar to the Equal Effects scenario with 5 phenotypes (Fig 1A). The relative improvement in accuracy provided by *mr.mash-rss* is, however, a little larger in this scenario because the method can borrow information across more phenotypes with equal effects. On the other hand, *SmvBayesC* “restrictive” did not take advantage from the larger number of phenotypes and provided a relative improvement over the univariate methods that was similar to that of the Equal Effects scenario with 5 phenotypes. We could not run the full version of *SmvBayesC* in this scenario because it was too computationally intensive.

## Applications

### Case study: Predicting blood cell traits in the UK Biobank

To evaluate *mr.mash-rss* on a real data application, we sought to predict blood cell traits from genotypes using the UK Biobank data. The UK Biobank is a dataset of roughly 500,000 individuals with genetic and phenotypic data [30]. We focused on a subset of 16 blood cell traits that have been used for quantitative genetic analyses in previous work [21]. After a series of filters (see S1 Text for details), our data consisted of *n* = 244, 049 individuals and *p* = 1, 054, 330 HapMap3 variants, as has been previously recommended [31]. The 244K individuals were split into 5 non-overlapping groups to perform 5-fold cross-validation. Each method was trained on the data from 4 groups and prediction accuracy was computed in the remaining fifth group. This procedure was repeated five times, once for each fold. Given that *SmvBayesC* is too computationally intensive for this many phenotypes, we compared *mr.mash-rss, LDpred2-auto*, and *SBayesR* in the real data application.

The results of this analysis are summarized in Fig 3 and S1 Table. Overall, the three methods performed similarly. This result is similar to what we found in the “Shared Effects in Subgroup” simulation scenario, which was designed to be reflective of the complex genetic architecture and effect sharing patterns of actual complex traits. However, *mr.mash-rss* was the most accurate for 14 out 16 blood cell phenotypes. The relative change in mean prediction accuracy compared to *LDpred2-auto* ranged from -0.6% (Eosinophil Percentage) to 32.8% (Basophill Percentage), with an average of 5.4% (Table 1). The relative change in mean prediction accuracy compared to *SBayesR* ranged from -1.9% (Eosinophil Percentage) to 13.9% (Basophill Percentage), with an average of 2.7%(Table 1). The better performance of *SBayesR* compared to *LDpred2-auto* may be due to a more flexible prior that can better approximate the actual distribution of the genetic effects.

**Table 1.** Percentage change in mean *R*^2^ of *mr.mash-rss* relative to *LDpred2-auto* and *SBayesR* for the 16 blood cell traits in the full and sampled UK Biobank data.

| Phenotype | Full data |  | Sampled data |  |
| --- | --- | --- | --- | --- |
|  | *LDpred2-auto* | *SBayesR* | *LDpred2-auto* | *SBayesR* |
| Red Blood Cell Counts (RBC#) | 2.6 | 1.1 | 18.1 | 6.1 |
| Haemoglobin Concentration (HGB) | 0.1 | <u>-1.6</u> | 13.5 | <u>-0.7</u> |
| Mean Corpuscular Volume (MCV) | 2.3 | 1.3 | 7.7 | 1.3 |
| Red Blood Cell Volume Distribution Width (RDW) | 4.8 | 3.3 | 19.6 | 7.5 |
| Mean Sphered Cell Volume (MSCV) | 6.1 | 4.4 | 17.2 | 7.6 |
| Reticulocyte Percentage (RET%) | 6.3 | 3.5 | 20.1 | 8.3 |
| High Light Scatter Reticulocytes Percentage (HLR%) | 4.0 | 1.4 | 16.3 | 4.7 |
| Platelet Count (PLT#) | 1.8 | 0.9 | 11.3 | 4.3 |
| Plateletcrit (PCT) | 1.6 | 0.6 | 10.9 | 3.0 |
| Platelet Distribution Width (PDW) | 1.7 | 0.7 | 9.0 | 1.7 |
| White Blood Cell Count (WBC#) | 3.2 | 1.8 | 18.3 | 4.3 |
| Lymphocyte Percentage (LYMPH%) | 6.9 | 5.1 | 32.9 | 11.5 |
| Monocyte Percentage (MONO%) | 3.2 | 1.3 | 7.2 | <u>-1.4</u> |
| Neutrophil Percentage (NEUT%) | 9.7 | 7.1 | 36.0 | 13.8 |
| Eosinophil Percentage (EO%) | <u>-0.6</u> | <u>-1.9</u> | 12.4 | <u>-2.4</u> |
| Basophil Percentage (BASO%) | 32.8 | 13.9 | 122.3 | 13.5 |

Underlined are the negative values, *i.e.*, those instances where *mr.mash-rss* produces lower accuracy than the competing method.

**Fig 3.**
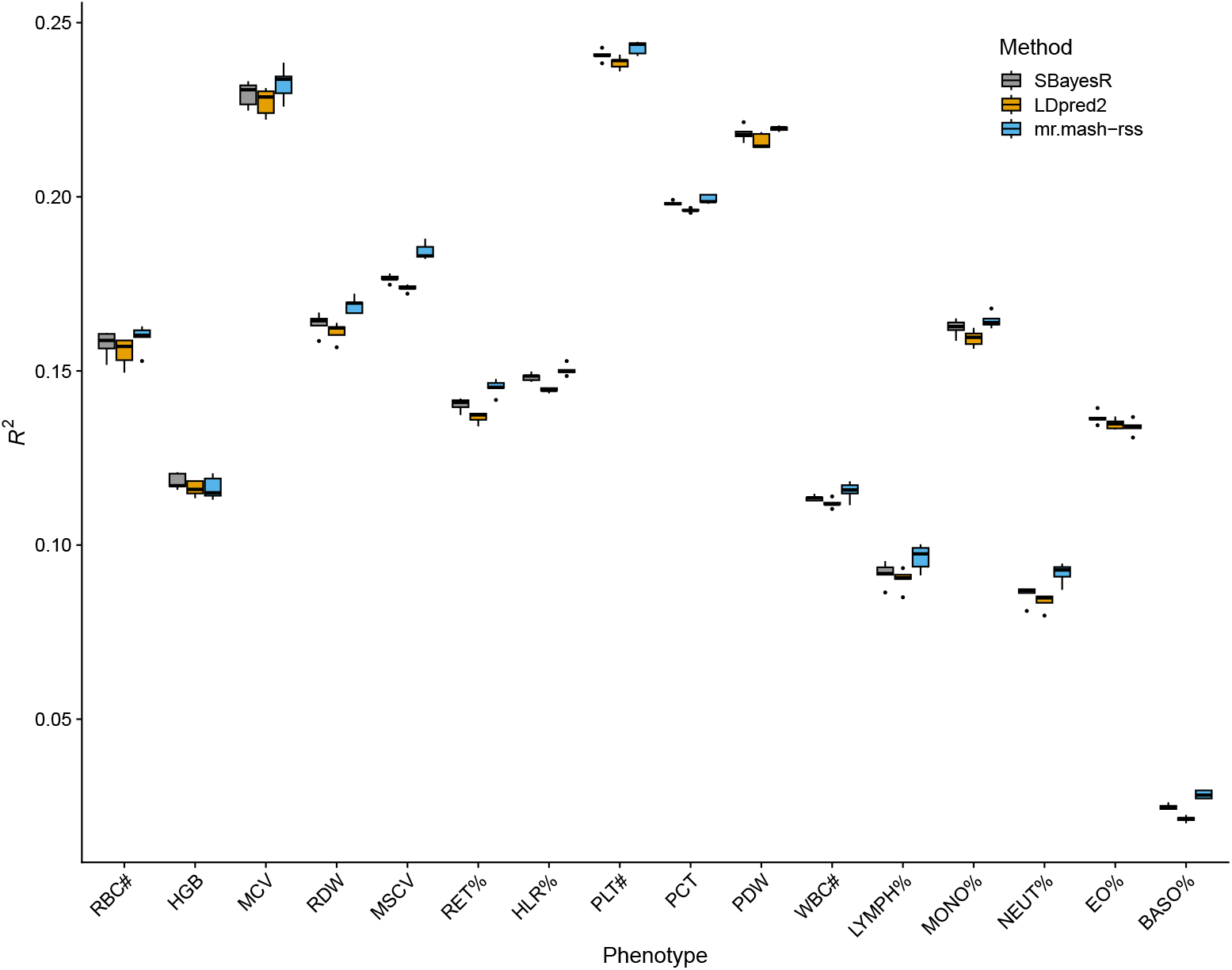
Prediction accuracy for the 16 blood cell traits in the full UK Biobank data. The thick, black line in each box gives the median *R*^2^.

In accordance with the simulations, the improvement in accuracy tended to be largest for phenotypes with lower genomic heritability (though this relationship is only suggestive) as shown in Fig 4. With lower signal-to-noise ratio, leveraging the sharing of effects in a multivariate analysis can give greatest improvements. This can be seen, for example, for Neutrophil Percentage (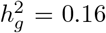; S2 Table), which has been shown to share putative causal variants with Lymphocyte Percentage (Fig. 3C in [21]) and is one of the phenotypes showing a greater improvement from using *mr.mash-rss*. On the other hand, the three platelet phenotypes have higher genomic heritability (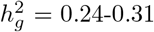; S2 Table) and despite some sharing of causal variants (Fig. 3C in [21]), the improvements in accuracy from using *mr.mash-rss* are very small.

**Fig 4.**
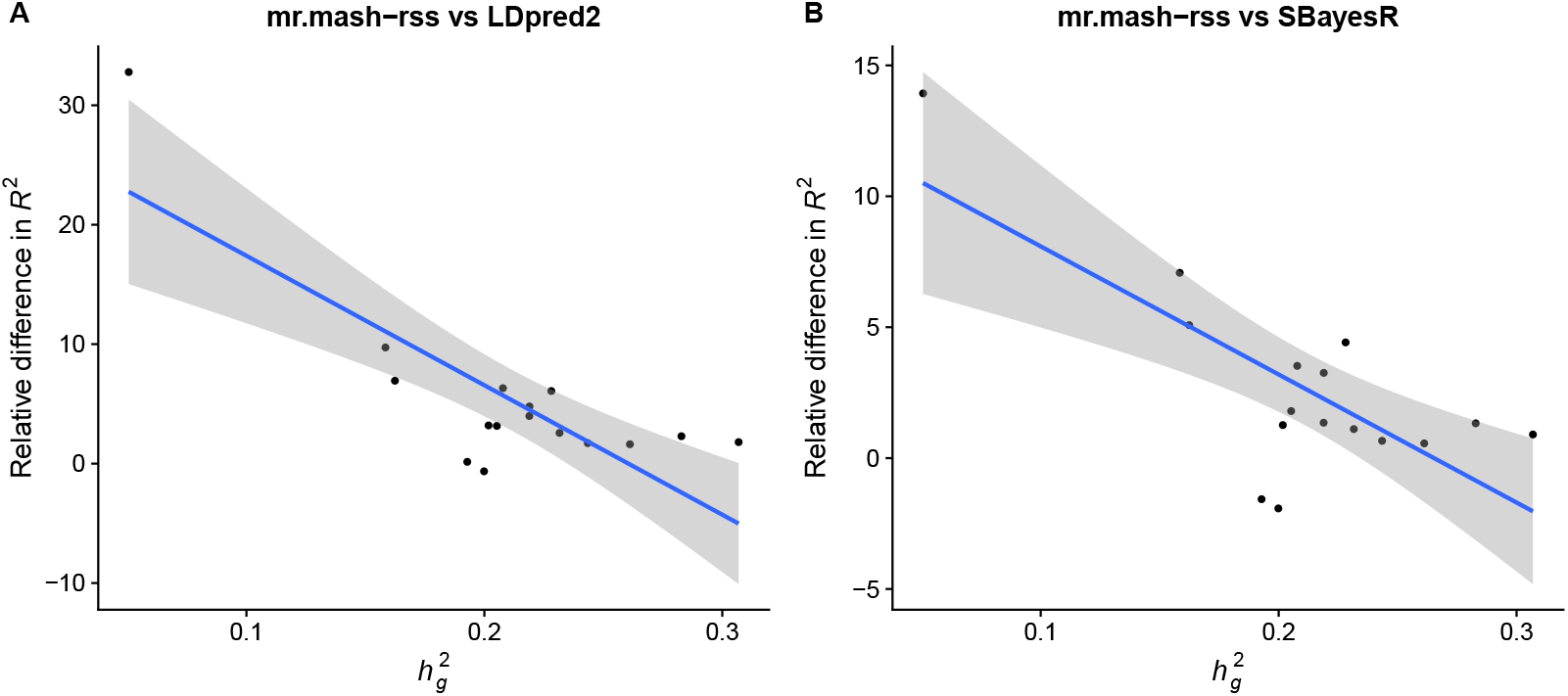
Relationship between improvement in prediction accuracy and genomic heritability in the full UK Biobank data. Phenotypes are plotted along the x-axis by their genomic heritability (*h*^2^) and along the y-axis by the change in *R*^2^ relative to the *LDpred2-auto* (Panel A) and *SBayesR* (Panel B); that is,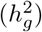 (*mr.mash-rss*) - *R*^2^(other method))/*R*^2^(other method). The blue line represents the linear regression fit with 95% confidence bands.

Previous analyses have shown that phenotypes with smaller sample size gain more advantage from multivariate modeling [18, 32]. We hypothesized that more substantial improvements in prediction accuracy from using *mr.mash-rss* could be obtained with a smaller sample size. Thus, we repeated the same analysis on 75,000 individuals, randomly sampled from the total of 244,000. The results, summarized in Fig 5 and and S1 Table, showed that this is indeed the case.

**Fig 5.**
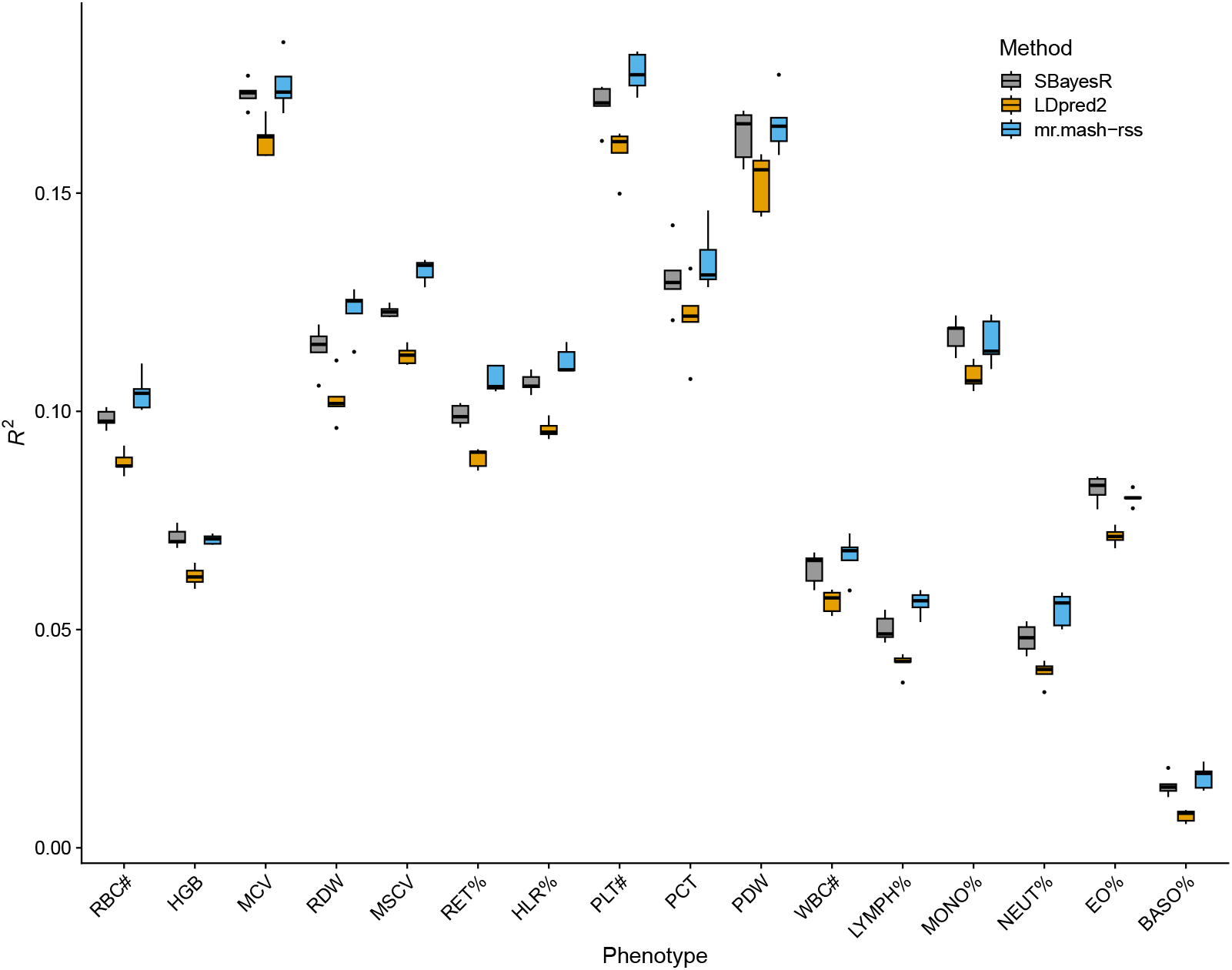
Prediction accuracy for the 16 blood cell traits in the sampled UK Biobank data. The thick, black line in each box gives the median *R*^2^.

In fact, the relative change in mean prediction accuracy compared to *LDpred2-auto* ranged from 7.2% (Monocyte Percentage) to 122.3% (Basophill Percentage), with an average of 23.3% (Table 1). This is about 4 times larger than the average relative change in mean prediction accuracy using the full data. The relative change in mean prediction accuracy compared to *SBayesR* ranged from -2.4% (Eosinophil Percentage) to 13.5% (Neutrophil Percentage), with an average of 5.2% (Table 1). This is about 2 times larger than the average relative change in mean prediction accuracy using the full data

## Discussion

In this work, we have introduced *mr.mash-rss*, the summary data version of a recently developed empirical Bayes multivariate multiple regression method [18]. Like *mr.mash, mr.mash-rss* enjoys (1) the ability to learn patterns of effect sharing across phenotypes; (2) the ability to model dozens of phenotypes jointly; (3) computational efficiency. However, *mr.mash-rss* addresses two important limitations of *mr.mash*— the need for individual-level data and the lack of scalability to biobank-size data.

Through an array of simulations and real data analysis using the UK Biobank, we showed that *mr.mash-rss* is competitive with state-of-the-art univariate and multivariate PGS methods. Of note, *mr.mash-rss* outperformed competing methods in 14 out of 16 blood cell phenotypes, although the magnitude of the improvement varied across phenotypes, from modest to substantial. This highlights that the general *mr.mash* model can adapt to either more sparse (e.g., for gene expression [18]) or more dense (e.g., for complex traits) genetic architectures. We also showed that the improvement in prediction accuracy from using *mr.mash-rss* increased substantially with a smaller sample size. This holds good promise for improving prediction accuracy for phenotypes that are difficult to measure and in samples of individuals of non-European descent, which are usually much smaller [33]. In addition, the performance of the *mr.mash* model depends on the accuracy of the “data-driven” covariance matrices [18]. Thus, advances in covariance matrix estimation can potentially lead to improvements in prediction accuracy.

A limitation of *mr.mash-rss* is that it requires the summary statistics to be computed on the same samples for each phenotype. In other words, there should not be missing data in ***Y*** in 1. Dealing with arbitrary patterns of missing data in multivariate models is not a trivial problem [34] and is an area where more research is needed. However, if individual-level data are available, missing values may be imputed before the prediction analysis. In fact, recent work has shown that imputing missing values results in improved prediction accuracy of PGS and power in GWAS [35, 36]. In addition, in specific cases such as with sample non-overlap across phenotypes, some simplifications arise that allow for models like *mr.mash-rss* to be fitted efficiently [37].

## Supporting information

**S1 Table Mean prediction** *R*^2^ **across folds for the 16 blood cell traits in the full and sampled UK Biobank data**.

**S2 Table Mean** 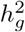 **across training sets for the 16 blood cell traits in the full UK Biobank data**.

**S1 Text. Detailed methods**. Detailed description of the methods, including: derivations of the *mr.mash-rss* algorithms, data preparation; simulations; methods compared; data analysis.

## Data and code availability

The genotype and phenotype data used in our analyses are available from UK Biobank (https://www.ukbiobank.ac.uk/). All code implementing the simulations and the compiled results generated from our simulations as well as all the code used to analyze the data are available at https://github.com/morgantelab/mr_mash_rss_analysis.

## Acknowledgments

This research was conducted using the UK Biobank Resource under application number 129216. We thank Gao Wang, Yuxin Zou, Peter Carbonetto and Matthew Stephens for useful discussions. Research reported in this publication was supported by the National Institute of General Medical Sciences of the National Institutes of Health under Award Number R35GM146868 to FM. The content is solely the responsibility of the authors and does not necessarily represent the official views of the National Institutes of Health. PS acknowledges support from Open Discovery Innovation Network (ODIN) under grant number NNF20SA0061466.

