## Supplementary material for "Improving polygenic prediction from summary data by learning patterns of effect sharing across multiple phenotypes": S1 Text

### Conventions used in mathematical expressions

For the mathematical expressions below, we denote matrices as bold, uppercase letters (e.g., $\boldsymbol{A}$), column vectors as bold, lowercase letters ($\boldsymbol{a}$), and scalars are written in plain, lowercase letters ($a$). We use $\mathbb{R}^{m\times n}$ to denote the set of real $m \times n$ matrices, we use $\boldsymbol{I}_n$ to denote the $n \times n$ identity matrix, $\mathbf{1}_n$ is a column-vector of ones of length $n$, and $\mathrm{tr}(\boldsymbol{A}) = \sum_{i=1}^{n} a_{ii}$ denotes the trace of $n \times n$ matrix $\boldsymbol{A}$.

For a matrix $\mathbf{A}$, let $\mathbf{A}_{\{i,j\}}$ denote its $i, j^{th}$ element, let $\mathbf{A}_{\{,j\}}$ and $\mathbf{A}_{\{j,\}}$ denote the $j^{th}$ column and $j^{th}$ row of $\mathbf{A}$, respectively. Further, let $\mathbf{A}_{\{,-j\}}$ denote the matrix with its $j^{th}$ column removed and let $\mathbf{A}_{\{-j,\}}$ denote the matrix with $j^{th}$ row removed. For a matrix of the form $\mathbf{A}^\intercal\mathbf{B}$ with $\mathbf{A}$ having $j^{th}$ column $\mathbf{a}_j$, the quantity $\mathbf{a}_j^\intercal\mathbf{B}$ can be obtained by subsetting as: $\mathbf{a}_j^\intercal\mathbf{B} = \mathbf{A}^\intercal\mathbf{B}_{\{j,\}}$ and $\mathbf{a}_j^\intercal\mathbf{a}_j = \mathbf{A}^\intercal\mathbf{A}_{\{j,j\}}$.

For describing Bayesian calculations, our convention is to use the "0" subscript to indicate priors and the "1" subscript to indicate posteriors.

### 1. *mr.mash* model

In *mr.mash* and *mr.mash-rss*, the response follows a Bayesian multivariate multiple regression model; that is,

$$\begin{aligned} \boldsymbol{Y} &= \boldsymbol{X}\boldsymbol{B} + \boldsymbol{E} \\ \boldsymbol{E} &\sim MN_{n\times r}\left(\mathbf{0}, \boldsymbol{I}_n, \boldsymbol{V}\right), \end{aligned} \tag{5}$$

where $\boldsymbol{Y}$ is an $n \times r$ matrix, $\boldsymbol{X}$ is an $n \times p$ matrix, $\boldsymbol{B}$ is a $p \times r$ matrix, and $\boldsymbol{V}$ is $r \times r$. The effect of the $j^{th}$ variable is denoted by the $r$-vector $\boldsymbol{b}_j$, where $\boldsymbol{b}_j^T$ is the $j^{th}$ row of $\boldsymbol{B}$. The prior on $\boldsymbol{B}$ is

$$\boldsymbol{b}_j \mid \boldsymbol{w}_0, \mathscr{S}_0 \sim \sum_{k=1}^{K} w_{0,k} N_r(\mathbf{0}, \boldsymbol{S}_{0,k}), \quad j = 1, \ldots, p. \tag{6}$$

### 2. Fitting the model with sufficient statistics

In the next section we describe methods for fitting the *mr.mash* model that rely on the data only through $\boldsymbol{X}^\intercal\boldsymbol{X}$, $\boldsymbol{X}^\intercal\boldsymbol{Y}$, and $\boldsymbol{Y}^\intercal\boldsymbol{Y}$, the sufficient statistics for the model in (5).

#### *2.1. Update of* $\mathbf{B}$.

We approximate the posterior distribution of $\boldsymbol{B}$ through variational inference, a method for posterior inference that approximates the posterior through optimization rather than sampling, as in Monte Carlo methods. A class of distributions is specified for the approximate posterior, denoted by $q(\boldsymbol{B})$. The optimal $q$ from within this class is found by minimizing the Kullback-Leibler divergence ($D_{KL}$) of the approximation from the true posterior or, equivalently, by maximizing the evidence lower bound (ELBO) with respect to $q$. The ELBO is defined to be

$$F(q; \boldsymbol{V}, \boldsymbol{w}_0) := \log(p(\boldsymbol{Y}|\boldsymbol{X}, \boldsymbol{V})) - D_{KL}\left(q(\boldsymbol{B}) \mid\mid p(\boldsymbol{B}|\boldsymbol{X}, \boldsymbol{Y}, \boldsymbol{V}, \boldsymbol{w}_0)\right) \tag{7}$$

where the first term is the log marginal likelihood of the model and the second term is the Kullback-Leibler divergence of the approximate posterior from the true posterior.

Following [7], in *mr.mash-rss*, we assume a mean-field approximation for $q(\boldsymbol{B})$; that is, we restrict the approximate posterior to take the form:

$$q(\boldsymbol{B}) = \prod_{j=1}^{p} q_j(\boldsymbol{b}_j), \tag{8}$$

where the $q_j$ is a probability distribution on the $j^{th}$ row of $\boldsymbol{B}$. Subject to this constraint, the ELBO can be expressed in a more tractable form:

$$F(q_1,\ldots,q_p;\boldsymbol{V},\boldsymbol{w}_0) = E_q\left[\log(p(\boldsymbol{Y}|\boldsymbol{X},\boldsymbol{V},\boldsymbol{B}))\right] + \sum_{j=1}^{p} D_{KL}\left(q_j(\boldsymbol{b}_j)||g(\boldsymbol{b}_j)\right) \tag{9}$$

where the expectation in the first term is with respect to the approximate posterior, $q$, and $g(\boldsymbol{b}_j)$ denotes the prior on $\boldsymbol{b}_j$.

We maximize $F(q;\boldsymbol{V},\boldsymbol{w}_0)$ sequentially with respect to each $q_j$, holding $q_{j'}$ constant for $j' \neq j$. It can be shown that the optimal $q_j$ is a $K$-component Gaussian mixture of the following form:

$$q_j(\boldsymbol{b}_j) = \sum_{k=1}^{K} w_{1,k}^{(j)} \mathrm{N}_r(\boldsymbol{b}_{1,k}^{(j)}, \mathbf{S}_{1,k}^{(j)}). \tag{10}$$

The parameters of the mixture in (10) are functions of the coefficients of a least-squares multivariate regression of $\bar{\boldsymbol{R}}_{(j)}$ on $\boldsymbol{x}_j$, where $\bar{\boldsymbol{R}}_{(j)}$ denotes the residuals with the effect of $\boldsymbol{b}_j$ removed; i.e, $\bar{\boldsymbol{R}}_{(j)}$ is the $n \times r$ matrix whose $i^{th}$ row is $\boldsymbol{Y}_{\{i,\}} - \sum_{h\neq j} x_{ih}\boldsymbol{b}_h^{\intercal}$. The mean, weight, and covariance for the $k^{th}$ component and $j^{th}$ predictor are given by the following expressions:

$$\begin{aligned}
\boldsymbol{S}_{1,k}^{(j)} &= \left(\boldsymbol{S}_{0,k}^{-1} + \widehat{\boldsymbol{S}}_j^{-1}\right)^{-1} \\
w_{1,k}^{(j)} &\propto w_{0,k} \frac{|\widehat{\boldsymbol{S}}_j|^{1/2}}{|\boldsymbol{S}_{0,k} + \widehat{\boldsymbol{S}}_j|^{1/2}} \exp\left(\frac{1}{2}\left(\boldsymbol{b}_{1,k}^{(j)}\right)^T \left(\boldsymbol{S}_{1,k}^{(j)}\right)^{-1} \boldsymbol{b}_{1,k}^{(j)}\right) \\
\boldsymbol{b}_{1,k}^{(j)} &= \boldsymbol{S}_{1,k}^{(j)} \widehat{\boldsymbol{S}}_j^{-1} \widehat{\boldsymbol{b}}_j,
\end{aligned} \tag{11}$$

where $\widehat{\boldsymbol{S}}_j = \boldsymbol{V}/\boldsymbol{x}_j^{\intercal}\boldsymbol{x}_j$ and $\widehat{\boldsymbol{b}}_j^{\intercal} = \boldsymbol{x}_j^{\intercal}\bar{\boldsymbol{R}}_{(j)}/\boldsymbol{x}_j^{\intercal}\boldsymbol{x}_j$. The quantity $\boldsymbol{x}_j^{\intercal}\bar{\boldsymbol{R}}_{(j)}$ is the $j^{th}$ row of $\boldsymbol{X}^{\intercal}\bar{\boldsymbol{R}}_{(j)}$, which is found as

$$\boldsymbol{X}^{\intercal}\bar{\boldsymbol{R}}_{(j)} = \boldsymbol{X}^{\intercal}\boldsymbol{X}\bar{\boldsymbol{B}} - \boldsymbol{X}^{\intercal}\boldsymbol{Y} + \boldsymbol{X}^{\intercal}\boldsymbol{x}_j\boldsymbol{b}_j^{\intercal}. \tag{12}$$

The quantity $\boldsymbol{X}^{\intercal}\boldsymbol{x}_j$ is the $j^{th}$ column of $\boldsymbol{X}^{\intercal}\boldsymbol{X}$ and can be found by subsetting: $\boldsymbol{X}^{\intercal}\boldsymbol{x}_j = \boldsymbol{X}^{\intercal}\boldsymbol{X}_{\{,j\}}$ and $\bar{\boldsymbol{B}}$ is the $p \times r$ matrix whose $j^{th}$ row is $\bar{\boldsymbol{b}}_j^{\intercal}$, defined below in (14).

We define the function BMSR-mix-ss to be one that returns the coefficients $\boldsymbol{b}_{1,k}^{(j)}$, $w_{1,k}^{(j)}$, and $\boldsymbol{S}_{1,k}^{(j)}$, $k = 1,\ldots,K$ of the distribution $q_j$, given $\boldsymbol{x}_j^{\intercal}\boldsymbol{x}_j$, $\boldsymbol{x}_j^{\intercal}\bar{\boldsymbol{R}}_{(j)}$, and the prior:

$$\text{BMSR-mix-ss}(\boldsymbol{x}_j^{\intercal}\boldsymbol{x}_j, \boldsymbol{x}_j^{\intercal}\bar{\boldsymbol{R}}_{(j)}, \boldsymbol{w}_0, \mathscr{S}_0) := \left(w_{1,1}^{(j)}\ldots, w_{1,K}^{(j)}, \boldsymbol{b}_{1,1}^{(j)}\ldots, \boldsymbol{b}_{1,K}^{(j)}, \boldsymbol{S}_{1,1}^{(j)}\ldots, \boldsymbol{S}_{1,K}^{(j)}\right) \tag{13}$$

Note: in the *mr.mash-rss* algorithm, the updates use BMSR-mix-ss$(\boldsymbol{x}_j^{\intercal}\boldsymbol{x}_j, \boldsymbol{x}_j^{\intercal}\bar{\boldsymbol{R}}_{(j)}, \boldsymbol{w}_0, \mathscr{S}_0)$ to iteratively update the $\boldsymbol{b}_j$ for $j = 1,\ldots,p$. If instead BMSR-mix-ss$(\boldsymbol{x}_j^{\intercal}\boldsymbol{x}_j, \boldsymbol{x}_j^{\intercal}\boldsymbol{Y}, \boldsymbol{w}_0, \mathscr{S}_0)$ is used, the estimates give the posterior distribution of $\boldsymbol{b}_j$ in the regression of $\boldsymbol{Y}$ on $\boldsymbol{x}_j$ under the prior in (6).

The parameters calculated in (13) are used to updated $\boldsymbol{b}_j$ by its posterior mean $\bar{\boldsymbol{b}}_j := E_{q_j}(\boldsymbol{b}_j)$. The update is given below as a function of the component means, covariances, and weights of the distribution in (10).

$$\begin{aligned}
\bar{\boldsymbol{b}}_j &= \sum_{k=1}^{K} w_{1,k}^{(j)}\boldsymbol{b}_{1,k}^{(j)} \\
\boldsymbol{S}_j &= \sum_{k=1}^{K} w_{1,k}^{(j)}\left(\boldsymbol{S}_{1,k}^{(j)} + \boldsymbol{b}_{1,k}^{(j)}\left(\boldsymbol{b}_{1,k}^{(j)}\right)^{\intercal}\right) - \bar{\boldsymbol{b}}_j\bar{\boldsymbol{b}}_j^{\intercal}
\end{aligned} \tag{14}$$

#### *2.2. Update of $\boldsymbol{w}_0$.*

The prior (6) is determined by the pre-specified covariance matrices, $\mathscr{S}_0$, and the prior weights given by $\boldsymbol{w}_0$. We assume that the elements of $\mathscr{S}_0$ are known, and we estimate $\boldsymbol{w}_0$ from the data by maximizing the ELBO. Its update, derived in [7], is

$$\widehat{w}_{0,k} = \frac{1}{p}\sum_{j=1}^{p} w_{1,k}^{(j)}, \quad k = 1,\ldots,K, \tag{15}$$

where the $w_{1,k}^{(j)}$, $j = 1, \ldots, p; k = 1, \ldots, K$ are obtained from BMSR-mix-ss defined in (13). We define the function UPDATE-WEIGHTS to be a function that outputs $\boldsymbol{w}_0$ according to the expressions (15) above.

$$\text{UPDATE-WEIGHTS}(\boldsymbol{W}) := (\hat{w}_{0,1}, \ldots, \hat{w}_{0,K}), \tag{16}$$

#### *2.3. Update of* $\mathbf{V}$.

The update of $\boldsymbol{V}$ is also accomplished by maximizing the ELBO with respect to $\boldsymbol{V}$. The update of [7] can be expressed in terms of sufficient statistics as follows:

$$\boldsymbol{V} = \text{ERSS}/n, \tag{17}$$

In the expression above, ERSS is equal the expected residual sum of squares, and it can be expressed using sufficient statistics as follows:

$$\begin{aligned} \text{ERSS} &= E_q\left(||\boldsymbol{Y} - \boldsymbol{X}\boldsymbol{B}||^2\right) \\ &= \bar{\boldsymbol{R}}^\intercal \bar{\boldsymbol{R}} + \sum_{j=1}^{p} \boldsymbol{x}_j^\intercal \boldsymbol{x}_j \boldsymbol{S}_j \end{aligned} \tag{18}$$

where

$$\begin{aligned} \bar{\boldsymbol{R}}^\intercal \bar{\boldsymbol{R}} &:= \left(\boldsymbol{Y} - \boldsymbol{X}\bar{\boldsymbol{B}}\right)^\intercal \left(\boldsymbol{Y} - \boldsymbol{X}\bar{\boldsymbol{B}}\right) \\ &= \boldsymbol{Y}^\intercal \boldsymbol{Y} - \bar{\boldsymbol{B}}^\intercal \boldsymbol{X}^\intercal \boldsymbol{Y} - \boldsymbol{Y}^\intercal \boldsymbol{X}\bar{\boldsymbol{B}} + \bar{\boldsymbol{B}}^\intercal \boldsymbol{X}^\intercal \boldsymbol{X}\bar{\boldsymbol{B}} \end{aligned} \tag{19}$$

We define the function UPDATE-RESID-COV-SS to be a function that outputs $\boldsymbol{V}$ according to the expressions (17) and (18) above.

$$\text{UPDATE-RESID-COV-SS}(\boldsymbol{X}^\intercal\boldsymbol{X}, \boldsymbol{X}^\intercal\boldsymbol{Y}, \boldsymbol{Y}^\intercal\boldsymbol{Y}, \boldsymbol{B}, \boldsymbol{S}_1, \ldots, \boldsymbol{S}_p) := \boldsymbol{V} \tag{20}$$

#### *2.4. Calculation of the ELBO*

From (9), the ELBO for the *mr.mash-rss* is

$$F(q_1, \ldots, q_p; \boldsymbol{V}, \boldsymbol{w}_0) = E_q\left[\log\left(p(\boldsymbol{Y} \mid \boldsymbol{X}, \boldsymbol{B}, \boldsymbol{V}, \boldsymbol{w}_0)\right] + \sum_{j=1}^{p} E_q\left[\log\left(\frac{p(\boldsymbol{b}_j)}{q_j(\boldsymbol{b}_j)}\right)\right], \tag{21}$$

with $p(\boldsymbol{b}_j)$ denoting the prior density of $\boldsymbol{b}_j$ and the second term representing the negative KL-divergence of $p$ from $q_j$. The first term in (21) can be expressed with sufficient statistics as :

$$E_q\left[\log\left(p(\boldsymbol{Y} \mid \boldsymbol{X}, \boldsymbol{B}, \boldsymbol{V}, \boldsymbol{w}_0)\right] = c - \frac{n}{2}\log|V| - \frac{1}{2}\text{tr}\left(\boldsymbol{V}^{-1}\bar{\boldsymbol{R}}^\intercal\bar{\boldsymbol{R}}\right) + \sum_{j=1}^{p}\text{tr}\left(\boldsymbol{V}^{-1}\boldsymbol{S}_j\right)\boldsymbol{x}_j^\intercal\boldsymbol{x}_j \tag{22}$$

where the $c$ represents terms that are constant with respect to $q$, $\boldsymbol{V}$, and $\boldsymbol{w}_0$ and $\bar{\boldsymbol{R}}^\intercal\bar{\boldsymbol{R}}$ is calculated using (19). The $j^{th}$ summand in the second term of (21) is

$$\begin{aligned} E_q\left[\log\left(\frac{q_j(\boldsymbol{b}_j)}{p(\boldsymbol{b}_j)}\right)\right] &= -D_{KL}(q_j(\boldsymbol{b}_j) \mid\mid p(\boldsymbol{b}_j)) \\ &= \log\left[\sum_{k=1}^{K} w_{0,k}\left(\frac{|\widehat{\boldsymbol{S}}_j|^{1/2}}{|\boldsymbol{S}_{0,k} + \widehat{\boldsymbol{S}}_j|^{1/2}}\right)\exp\left(-\frac{1}{2}\left(\boldsymbol{b}_{1,k}^{(j)}\right)^\intercal\left(\boldsymbol{S}_{1,k}^{(j)}\right)^{-1}\boldsymbol{b}_{1,k}^{(j)}\right)\right] + \\ &\quad \frac{1}{2}\left[\text{tr}(\boldsymbol{V}^{-1}\left(-\bar{\boldsymbol{b}}_j\boldsymbol{x}_j^\intercal\bar{\boldsymbol{R}}_j - \bar{\boldsymbol{R}}_j^\intercal\boldsymbol{x}_j\bar{\boldsymbol{b}}_j^\intercal + (\boldsymbol{S}_j + \bar{\boldsymbol{b}}_j\bar{\boldsymbol{b}}_j^\intercal)\boldsymbol{x}_j^\intercal\boldsymbol{x}_j\right)\right] \end{aligned} \tag{23}$$

**Algorithm 1** *mr.mash*-sufficient

**Require:** $\boldsymbol{X}^\intercal\boldsymbol{X}$, $\boldsymbol{X}^\intercal\boldsymbol{Y}$, and $\boldsymbol{Y}^\intercal\boldsymbol{Y}$.
**Require:** Set of $K$ covariance matrices, $\mathscr{S}_0$.
**Require:** Initial estimates of the posterior mean coefficients, stored as a $p \times r$ matrix, $\bar{\boldsymbol{B}}^0$ whose $j^{th}$ row is $\boldsymbol{b}_j^\intercal$.
**Require:** Initial estimates of the prior mixture weights, $\boldsymbol{w}_0^0 = (w_{0,1}, \ldots, w_{0,K})$, and the $r \times r$ residual covariance matrix $\boldsymbol{V}^0$.
**Require:** Convergence threshold, tol $\geq 0$, and an upper limit on the number of iterations, $t_{\max}$.

```
t ← 0
δ ← ∞
ELBO^(0) ← F(q; V^0, w_0^0)
while δ > tol and t < t_max do
    t ← t + 1
    Compute X^T R̄ ← X^T Y − X^T X B̄
    Initialize W to a p × K matrix of zeros
    for j in 1, ..., p do
        Remove variable j from "expected residuals", X^T R̄_(j) ← X^T R̄ + X^T x_j b̄_j^T
        (w_{1,1}^(j), ..., w_{1,K}^(j), b_{1,1}^(j), ..., b_{1,K}^(j), S_{1,1}^(j), ..., S_{1,K}^(j)) ← BMSR-mix-ss(x_j^T x_j, x_j^T R̄_(j), V, 𝒮_0, w_0)   ▷ See (13).
        Compute posterior mean, b̄_j = Σ_{k=1}^K w_{1,k}^(j) b_{1,k}^(j)
        Compute posterior covariance, S_j = Σ_{k=1}^K w_{1,k}^(j) [ b_{1,k}^(j) (b_{1,k}^(j))^T + S_{1,k}^(j) ] − b̄_j b̄_j^T
        Store w_{1,1}^(j), ..., w_{1,K}^(j) in the j^th row of W
        Include variable j in "expected residuals", X^T R̄ ← X^T R̄_(j) − X^T x_j b̄_j^T
    Update prior weights, w_0 ← UPDATE-WEIGHTS(W)   ▷ See (16).
    Update residual covariance, V ← UPDATE-RESID-COV-SS(X^T X, X^T Y, Y^T Y, B̄, S_1, ..., S_p)   ▷ See (20).
    ELBO^(t) ← F(q; V, w_0)
    δ ← ELBO^(t) − ELBO^(t−1)
return B̄, V, w_0, ELBO^(t)
```

### 3. *mr.mash-rss* algorithm

Algorithms 1 and 2 provide pseudo-code to describe the fitting of the *mr.mash-rss* model. Algorithm 1 describes the variational algorithm to update the parameters $\boldsymbol{B}, \boldsymbol{V}$, and $\boldsymbol{w}_0$ using sufficient statistics.

The Algorithm 2 describes the *mr.mash-rss* algorithm, which takes as input the summary statistics and includes calculation of sufficient statistics and subsequent estimation of parameters via Algorithm 1. The summary statistics required, which are obtained from univariate simple linear regression models, are defined below.

- $\widehat{\boldsymbol{B}}$ is the $p \times r$ matrix whose $j, s^{th}$ element is the least-squares estimate of the slope of the regression of $\boldsymbol{y}_s$ on $\boldsymbol{x}_j$. Its $j^{th}$ row is denoted by $\widehat{\boldsymbol{b}}_j^\intercal$.

- $\widehat{\boldsymbol{\Sigma}}_{\widehat{b}}$ is the $p \times r$ matrix whose $j, s^{th}$ element is the estimated standard error of the estimated slope of the regression of $\boldsymbol{y}_s$ on $\boldsymbol{x}_j$. Its $j^{th}$ row is denoted by $\widehat{\boldsymbol{\sigma}}_{\widehat{b},j}^\intercal$.

- $\widehat{\boldsymbol{\Gamma}}$ is the $p \times p$ sample correlation matrix of $\boldsymbol{X}$.

The algorithm also requires $\boldsymbol{Y}^\intercal\boldsymbol{Y}$ (if available) and $n$.

†Operations inside this loop are elementwise multiplication and division.
‡This step calculates the PVE-adjusted z score. See [16] for details.

**Algorithm 2** *mr.mash-rss*

**Require:** $\widehat{\boldsymbol{B}}$, $\widehat{\boldsymbol{\Sigma}}_{\widehat{b}}$, $\widehat{\boldsymbol{\Gamma}}$, $\boldsymbol{Y}^\intercal\boldsymbol{Y}$, $n$. Let $\boldsymbol{v}_Y = (\boldsymbol{y}_1^\intercal\boldsymbol{y}_1, \ldots, \boldsymbol{y}_r^\intercal\boldsymbol{y}_r)/(n-1)$ denote the $r-$vector containing the sample variances of each response.

**Require:** Set of $K$ covariance matrices, $\mathscr{S}_0$.

**Require:** Initial estimates of the posterior mean coefficients, stored as a $p \times r$ matrix, $\bar{\boldsymbol{B}}^0$ whose $j^{th}$ row is $\boldsymbol{b}_j^\intercal$.

**Require:** Initial estimates of the prior mixture weights, $\boldsymbol{w}_0^0 = (w_{0,1}, \ldots, w_{0,K})$, and the $r \times r$ residual covariance matrix $\boldsymbol{V}^0$.

**Require:** Convergence threshold, tol $\geq 0$, and an upper limit on the number of iterations, $t_{\max}$.

**for** $j$ in $1, \ldots, p$ **do**†

  $\widehat{\boldsymbol{z}}_j := \widehat{\boldsymbol{b}}_j / \widehat{\boldsymbol{\sigma}}_{\widehat{b},j}$

  $\boldsymbol{a}_j := (n-1)/(n-2+\widehat{\boldsymbol{z}}_j^2)$

  $\tilde{\boldsymbol{z}}_j := \sqrt{\boldsymbol{a}_j}\widehat{\boldsymbol{z}}_j$ ‡

  $d_j := mean((\boldsymbol{v}_Y \boldsymbol{a}_j / \widehat{\boldsymbol{s}}_j^2))$

  $\boldsymbol{x}_j^\intercal\boldsymbol{Y} = \sqrt{\boldsymbol{a}_j}\tilde{\boldsymbol{z}}_j \boldsymbol{v}_y / \widehat{\boldsymbol{s}}_j$

$\boldsymbol{D}_x := \mathrm{diag}(d_1, \ldots, d_p)$

$\boldsymbol{X}^\intercal\boldsymbol{X} = \boldsymbol{D}_x^{1/2}\widehat{\boldsymbol{\Gamma}}\boldsymbol{D}_x^{1/2}$

$$\boldsymbol{X}^\intercal\boldsymbol{Y} = \begin{bmatrix} \boldsymbol{x}_1^\intercal\boldsymbol{Y} \\ \vdots \\ \boldsymbol{x}_p^\intercal\boldsymbol{Y} \end{bmatrix}$$

  **return** $\boldsymbol{X}^\intercal\boldsymbol{X}$, $\boldsymbol{X}^\intercal\boldsymbol{Y}$, $\boldsymbol{Y}^\intercal\boldsymbol{Y}$.

$\bar{\boldsymbol{B}}$, $\boldsymbol{V}$, $\boldsymbol{w}_0$, ELBO$^{(t)}$ $\leftarrow$ mr.mash-sufficient($\boldsymbol{X}^\intercal\boldsymbol{X}$, $\boldsymbol{X}^\intercal\boldsymbol{Y}$, $\boldsymbol{Y}^\intercal\boldsymbol{Y}$, $n$, $\mathscr{S}_0$, $\boldsymbol{w}_0$, $\boldsymbol{V}$, $\bar{\boldsymbol{B}}$, tol, $t_{max}$). See Algorithm 1.

### 4. Details of the simulations

#### *4.1. Data preparation*

For the simulation analyses, we used real genotypes from the array data from the UK Biobank [2]. We kept individuals of Caucasian ancestry that self identified as White British. We computed relationships for every pairs of individuals based on genetic variants with minor allele frequency (MAF) greater than 0.01, minor allele count (MAC) greater than 5, Hardy-Weinberg equilibrium (HWE) test p-value greater than $10^{-10}$, and missing genotype rate smaller than 0.1. We retained individuals such that no pair had relationship coefficient greater than 0.025, for a total of 251,288 individuals. We randomly sampled 105,000 individuals and filtered out genetic variants that did not pass the same filters as above in this subset. These steps were performed using PLINK (v. 1.90b7) [3]. Our final dataset for subsequent analyses included 595,071 genetic variants. Missing genotypes were imputed with the mean genotype for the respective genetic variant.

#### *4.2. Simulation of phenotypes*

For each replicate, we simulated the $n \times r$ matrix of phenotypes for $n = 105,000$ and $r = 5$ (or 10 in one simulation scenario) following the multivariate regression model in (5). The predictor matrix $\boldsymbol{X}$ was the filtered genotype data from the previous step (that was centered so that its columns have means equal to 0) and the effect matrix $\boldsymbol{B}$ was chosen to obtain the desired pattern of effect sharing. In particular, the effect, $\boldsymbol{b}_j$, of each causal variant across phenotypes was sampled from a distribution that depends on the scenario:

- "Equal Effects", "Low $h_g^2$", "High Polygenicity", "More Phenotypes", where each causal variant affects all the phenotypes and has the same effect across phenotypes. $\boldsymbol{b}_j \sim N_r(\boldsymbol{0}, \boldsymbol{1}_r\boldsymbol{1}_r^T)$, where $\boldsymbol{1}_r = (1, \ldots, 1)^T \in \mathbb{R}^r$.
- "Mostly Null", where the causal variants affect only the first phenotype while the remaining

phenotypes are affected only by an non-genetic component. $\boldsymbol{b}_j \sim N_r(\boldsymbol{0}, \boldsymbol{S})$, where $\boldsymbol{S}$ is an $r \times r$ matrix of all zeros except for a single one, $s_{11} = 1$.

- "Shared Effects in Subgroups", where the effect of each causal variant is drawn such that it is equally likely to be shared (but not be equal) in phenotypes 1 through 3 or to be shared (but not be equal) in phenotypes 4 and 5. $\boldsymbol{b}_j \sim 0.5 \times N_r(\boldsymbol{0}, \boldsymbol{S}_1) + 0.5 \times N_r(\boldsymbol{0}, \boldsymbol{S}_2)$ where $\boldsymbol{S}_1$ is an $r \times r$ matrix with diagonal elements equal to 1 and off-diagonal elements equal to 0.9, and $\boldsymbol{S}_2$ is an $r \times r$ matrix with diagonal elements equal to 1 and off-diagonal elements equal to 0.7.

The effects of the non-causal variants were set to 0. The residual covariance $\boldsymbol{V}$ was a diagonal matrix where the diagonal elements were chosen to obtain the desired genomic heritability. This implies that the residuals were uncorrelated across phenotypes. In the "Mostly Null" scenario, the diagonal elements for the phenotypes unaffected by genotypes were set to be equal to 1.

Finally, we randomly split the data (*i.e.*, $\boldsymbol{X}$ and $\boldsymbol{Y}$) into a training set (including 100,000 individuals) and a test set (including 5,000 individuals). The simulation procedure was repeated 20 times to obtain 20 replicates.

#### 4.3. Computation of the summary statistics and LD matrices

We computed the summary statistics using only individuals in the training set by performing a univariate linear regression of each quantile normalized phenotype on each genetic variant, one at a time, using the `big_univLinReg` function from the R package bigstatsr (v.1.5.12) [8]. The effect sizes and their standard errors were used for subsequent analyses.

The linkage disequilibrium matrices were computed using 146,288 nominally unrelated individuals ($r_a < 0.025$ between any pair) individuals of European ancestry (*i.e.*, Caucasian and white British fields), that did not overlap with the 105,000 individuals used to compute the summary statistics and assess prediction accuracy. This procedure resulted in a set of "out-of-sample" LD matrices. We used the function `snp_cor` from the R package bigsnpr (v.1.12.2) [8] to compute correlations between variants for each chromosome separately, setting the correlation between variants that are 3 cM or more apart to be 0, as previously recommended [9].

#### 4.4. Computation of genetic predictors

We compared five different methods for polygenic prediction:

- *LDpred2-auto* [9]. This is a univariate Bayesian linear regression model fit by Gibbs sampling, where the prior on the effect size of each genetic variant is:

$$b_j \sim (1-\pi)\delta_0 + \pi N(0, \frac{h_g^2}{p}) \tag{24}$$

  We used the implementation in the R package bigsnpr (v.1.12.2) of this model. In particular, we used the `snp_ldpred2_auto` function that allows for the estimation of the hyperparameters (*i.e.*, $\pi$ and $h_g^2$) from the data. We ran *LDpred2-auto* 30 times with a sequence of 30 values equally spaced on a log scale from $10^{-4}$ to 1 as initial values of $\pi$, and an estimate from LD Score regression (*LDSC*; obtained using the function `snp_ldsc`) as initial value of $h_g^2$. In each run, we set the number of burn-in iterations to 500, the number of iterations (after burn-in) to 1,000, and kept the other arguments as their default values. We obtained a final estimate of the effect size for each variant as the average of the effect sizes from the 30 runs, after discarding the chains that diverged, as recommended by the *LDpred2-auto* authors [9].

- *SBayesR* [6]. This is a univariate Bayesian linear regression model fit by Gibbs sampling, where the prior on the effect size of each genetic variant is:

$$b_j \sim \pi_0\delta_0 + \sum_{k=1}^{K} \pi_k N(0, \gamma_k \sigma_b^2) \tag{25}$$

Here, we chose $K = 3$ and $\boldsymbol{\gamma} = (0.01, 0.1, 1.0)$. We used the implementation in the R package qgg (v. 1.1.2) [12] of this model. In particular, we used the `sblr` function that allows for the estimation of the hyperparameters (*i.e.*, $\boldsymbol{\pi}$ and $\sigma_b^2$) from the data. We initialized $h_g^2 = 0.1$ (which is used internally to initialize $\sigma_b^2$), we set the number of burn-in iterations to 1,000, the number of iterations to 5,000, the thinning parameter to 5, and kept the other arguments as their default values.

- *SmvBayesC* [4]. This is a multivariate Bayesian linear regression model fit by Gibbs sampling, where the prior on the effect size of each genetic variant is:

$$\begin{aligned} \boldsymbol{b}_j &= \boldsymbol{D}_j \boldsymbol{\beta}_j \\ \boldsymbol{\beta}_j &\sim N_r(\boldsymbol{0}, \boldsymbol{G}) \end{aligned} \tag{26}$$

where:

$$\boldsymbol{D} = \begin{bmatrix} d_{j1} & & \\ & \ddots & \\ & & d_{jr} \end{bmatrix}; \boldsymbol{G} = \begin{bmatrix} \sigma^2_{\beta 1} & \cdots & \sigma_{\beta 1 r} \\ \vdots & \ddots & \vdots \\ \sigma_{\beta 1 r} & \cdots & \sigma^2_{\beta r} \end{bmatrix}$$

$d_{js} \in 0, 1$ indicates whether variant $j$ has an effect on phenotype $s$. Each potential configuration of effect presence/absence across phenotypes – $\boldsymbol{d}_{jl}$ for $l = 1, 2, \ldots, L = 2^r$ – is assigned a prior probability $\Pi_l$ (subject to $\sum_{l=1}^{L} \Pi_l = 1$). $\boldsymbol{\Pi} = (\Pi_1, \Pi_2, \ldots, \Pi_L) \sim Dir(\boldsymbol{\alpha})$. We used the implementation in the R package qgg (v. 1.1.2) of this model. In particular, we used the `mtsblr` function that allows for the estimation of the hyperparameters (*i.e.*, $\boldsymbol{D}$ and $\boldsymbol{G}$) from the data. We initialized $h_g^2 = 0.1$ (which is used internally to initialize $\boldsymbol{G}$) and $\pi$ – the probability of a genetic variant having an effect on at least one phenotype – as 0.0001 in every scenario, except in "High Polygenicity" where it was 0.01. We set the number of burn-in iterations to 1,000, the number of iterations to 5,000, the thinning parameter to 5, and kept the other arguments as their default values.

- *SmvBayesC*-rest [5]. This is a multivariate Bayesian linear regression model similar to *SmvBayesC*, with the difference being that only 2 configurations for $\boldsymbol{d}_{jl}$ are allowed: an effect is either present in all or none of the phenotypes. We used the implementation in the R package qgg (v. 1.1.2) of this model. In particular, we used the `mtsblr` function that allows for the estimation of the hyperparameters (*i.e.*, $\boldsymbol{D}$ and $\boldsymbol{G}$) from the data. We initialized $h_g^2 = 0.1$ (which is used internally to initialize $\boldsymbol{G}$) and $\pi$ – the probability of a genetic variant having an effect on all the phenotypes – as 0.0001 in every scenario, except in "High Polygenicity" where it was 0.01. We set the number of burn-in iterations to 1,000, the number of iterations to 5,000, the thinning parameter to 5, and kept the other arguments as their default values.

- *mr.mash-rss*. This is the method introduced in this paper and described extensively above and the main text. We used the function `mr.mash.rss` implemented in the R package mr.mash.alpha (v. 0.3.14). We initialized the posterior mean of the regression coefficients, $\bar{\boldsymbol{B}}$, to 0. We computed an initial estimate of the residual covariance, $\boldsymbol{V}$, from summary statistics by following a similar approach to [17]. For each chromosome, we selected genetic variants whose $z$-score was less than 2 (in absolute value) in every trait. We computed an initial estimate of $\boldsymbol{V}$ using the selected genetic variants for all chromosomes as:

$$\boldsymbol{V} = \frac{1}{J} \sum_{j=1}^{J} \boldsymbol{z}_j \boldsymbol{z}_j^{\intercal} \tag{27}$$

where $J$ is the number of selected variants and $\boldsymbol{z}$ is an $r$-vector of $z$-scores across phenotypes. The mixture weights, $\boldsymbol{w}_0$, were initialized as 0.99 on $w_{0,0}$ (*i.e.*, the weight for the null component) and 0.01 split equally among the other mixture components. Both $\boldsymbol{V}$ and $\boldsymbol{w}_0$ were updated in the *mr.mash-rss* model fitting. $\boldsymbol{Y}^{\intercal}\boldsymbol{Y}$ was computed from the individual-level phenotypic data using only training individuals after quantile normalization. The algorithm was run until the difference in ELBO between two successive iterations was smaller than 0.01. To reduce runtime, at each iteration after the first 15 iterations, we dropped mixture components with estimated weight smaller than $10^{-8}$. Following [7], we used an expanded version of the mixture prior:

$$\boldsymbol{b}_j \mid \boldsymbol{w}_0, \boldsymbol{\omega}, \mathscr{U}_0 \sim w_{0,0}\delta_{\mathbf{0}} + \sum_{l=1}^{L}\sum_{t=1}^{T} w_{0,l,t} N_r(\mathbf{0}, \omega_l^2 \boldsymbol{U}_{0,t}), \tag{28}$$

where $\delta_{\mathbf{0}}$ is the delta mass function at zero, $\boldsymbol{\omega}$ is a vector scaling factors meant to capture the magnitude of the effect sizes and chosen as described in [13], and $\mathscr{U}_0$ is a list of normalized (such that the largest diagonal element was 1) covariance matrices meant to capture the patterns of effect sharing and specificity across phenotypes. The covariance matrices included "canonical" matrices:

- The identity matrix, $\boldsymbol{I}_r$.
- A matrix of all ones, $\boldsymbol{A} = \mathbf{1}_r\mathbf{1}_r^\intercal$, where $\mathbf{1}_r = (1, \ldots, 1)^\intercal$.
- $r$ rank-1 matrices of the form, $\boldsymbol{C}_s = \boldsymbol{c}_s\boldsymbol{c}_s^\intercal$, where $\boldsymbol{c}_s$ in an $r$-vector of all zeros except for a 1 at position $s$.
- Three matrices with diagonal elements equal to 1 and off-diagonal elements equal to $\sigma$, with $\sigma = 0.25, 0.5, 0.75$, respectively.

However, we also used "data-driven" matrices computed using the summary statistics for strong signals. In particular, for each chromosome, we selected genetic variants with $z$-score greater than 3 (in absolute value) in at least one phenotype. Then, we combined the selected genetic variants across chromosomes ($m$) to form the $m \times r$ matrix $\boldsymbol{Z}$.

- Three rank-1 matrices based on the top 3 principal components of $\boldsymbol{Z}$, such that $\boldsymbol{P}_e = \boldsymbol{v}_e\boldsymbol{v}_e^\intercal$ where $\boldsymbol{v}_e$ is the $e^{th}$ right singular vector, for $e = 1, 2, 3$.
- A rank-3 matrix based on the linear combination of the top 3 principal components of $\boldsymbol{Z}$, such that $\boldsymbol{P} = \frac{1}{m}\sum_{e=1}^{3}\sigma_e^2\boldsymbol{v}_e\boldsymbol{v}_e^\intercal$ where $\sigma_e^2$ is the $e^{th}$ squared singular value.
- A matrix based on the Empirical Bayes Matrix Factorization (EBMF) of $\boldsymbol{Z}$ as implemented in the R package flashier (v. 0.2.34). This is a matrix factorization method that produces a sparse low-rank approximation of the original matrix, while automatically selecting the rank of the approximation in a data-adaptive way [15]. $\boldsymbol{Q} = \frac{1}{m}\boldsymbol{F}\boldsymbol{L}^\intercal\boldsymbol{L}\boldsymbol{F}^\intercal$ where $\boldsymbol{L}$ is a matrix of estimated loadings and $\boldsymbol{L}$ is a matrix of estimated factors.
- A number of matrices such that $\boldsymbol{Q}_e = \boldsymbol{f}_e\boldsymbol{f}_e^\intercal$ where $f_e$ is the $e^{th}$ factor estimated by EBMF.

We then estimated the covariance matrices to use in *mr.mash-rss* by applying Extreme Deconvolution (ED) – implemented in the `cov_ed` function of the R package mashr (v. 0.2.71) – using the list of matrices above as initial estimates. In brief, this is an EM algorithm for fitting mixture models such as (6) and outputs a list of "denoised" covariance matrices [1]. This approach to estimate covariance matrices was introduced in [13] and has been used successfully in a few studies [7, 17].

Each method was applied to each chromosome separately. Phenotypic values for the test set individuals for a given phenotype $s$ were predicted as:

$$\hat{\boldsymbol{y}}_{\text{test},s} = \sum_{g=1}^{22} \boldsymbol{X}_{\text{test},g}\hat{\boldsymbol{b}}_{g,s} \tag{29}$$

where $\boldsymbol{X}_{\text{test},g}$ is the genotype matrix including only genetic variants on the $g^{th}$ chromosome, and $\hat{\boldsymbol{b}}_{g,s}$ is a vector of estimated effect sizes of the genetic variants on the $g^{th}$ chromosome on the $s^{th}$ phenotype.

Prediction accuracy was evaluated as $R^2$ from the regression of $\boldsymbol{y}_{\text{test},s}$ on $\hat{\boldsymbol{y}}_{\text{test},s}$, where $\boldsymbol{y}_{\text{test},s}$ is a vector of actual (quantile normalized) phenotypic values for the test set individuals.

### 5. Details of the real data application

#### 5.1. Data preparation

For the real data application, we used 16 blood cell phenotypes from UK Biobank, that have been used in previous genetic analyses [14, 17]. As in [17], we focused on a subset of the individuals that met the following criteria:

- Were identified as White British by field 22009 in UK Biobank.
- No mismatch between self-reported and genetic sex.
- Were not outliers for missing genotype rate and/or heterozygosity as identified by field 22027 in UK Biobank.
- Did not have close relative as identified by field 22021 in UK Biobank.
- Were not pregnant.
- Did not have blood related disease based on ICD10 codes.

Finally, after quantile normalization of each phenotype, we calculated the Mahalanobis distance for each individual ($\boldsymbol{y}_i^\intercal \hat{\Sigma}^{-1} \boldsymbol{y}_i$, where $\boldsymbol{y}_i$ is an $r$-vector of phenotypes for the $i^{th}$ individual and $\hat{\Sigma}$ is the sample phenotypic covariance matrix). We then excluded individuals falling above the 0.99 quantile of the $\chi^2_{16}$ distribution. After all the filtering, the final dataset included of $n = 244,049$.

For the majority of the analyses, we used a set of $p = 1,054,330$ HapMap3 variants from the imputed UK Biobank genotype data. This set has been recommended in previous prediction analyses because the genetic variants included provide a good coverage of the genome and are well imputed [10].

Finally, we randomly assigned each individual to one of five subsets to perform 5-fold cross-validation. This consists of training each model on 4 out of the 5 subsets data (*i.e.*, the training set) and evaluating the model performance on the remaining subset (*i.e.*, the test set). This procedure is repeated 5 times, once for each subset.

#### 5.2. Computation of the summary statistics and LD matrices

To compute summary statistics, we first adjusted each phenotype for the effect of sex, assessment center, age at recruitment, age $\times$ age, genotype measurement batch, and the first 10 genetic principal components by linear regression. The residuals for each phenotype were then quantile normalized and used as the response variable in a GWA analysis as described in Sec. 4.3. This procedure was performed 5 times, once for each training set in our 5-fold cross-validation scheme.

We computed a set of sparse per-chromosome LD matrices for each training set, using the same software and parameters as in Sec. 4.3. This procedure resulted in 5 sets of "in-sample" LD matrices.

#### 5.3. Computation of genetic predictors

We applied three different for polygenic prediction:

- *LDpred2-auto*. We used the same software and parameters as described in Sec. 4.4.
- *SBayesR*. We used the same software and parameters as described in Sec. 4.4.
- *mr.mash-rss*. We used the same software and a similar strategy to that described in Sec. 4.4, except for a few differences. First, we initialized the estimates of the regression coefficients with the estimates from *SBayesR*. *mr.mash-rss* solves a non-convex optimization problem and we found that a carefully chosen initialization could improve prediction accuracy [7]. Second, we computed the residual covariance in the same way as described above. However, we did not update it within the *mr.mash-rss* algorithm as that is the only step that requires an estimate of $\boldsymbol{Y}^\intercal \boldsymbol{Y}$, which might not always be available. Thus, we wanted to compare the methods in a realistic scenario. Third, we used a different strategy to select strong signals to compute the "data-driven" covariance matrices. For each training set:

– We performed association analyses for each trait and all biallelic autosomal SNPs with MAF greater than 0.001, and INFO score greater than 0.6 using the procedure described in Sec. 5.2.

– We obtained a list of 975 non-overlapping genomic regions that were used for fine-mapping the same 16 blood cell phenotypes in a previous study [17].

– For each of the 975 genomic regions, we fine-mapped the association results using the *SuSiE-RSS* method (implemented in the R package susieR (v. 0.12.40), with default parameters). For each phenotype, we selected genetic variants with the highest posterior inclusion probability (PIP) within each credible set (CS).

– We combined the lists of selected genetic variants across phenotypes and genomic regions to obtain a final list of genetic with strong evidence of association. These genetic variants were then used to compute the "data-driven" covariance matrices as described in Sec. 4.4.

#### *5.4. Analysis with sampled individuals*

We randomly sampled 15,000 individuals for each fold to obtain a total of 75,000 individuals. We repeated all the analyses described above, except that we did not recompute LD matrices and used the ones already computed for the full data analysis. Given the much smaller sample size, we also increased the MAF threshold to 0.01 for the association analysis used as part of the prior computation procedure.

### 6. Computing environment

All the analyses were run on Linux machines (Rocky Linux 8.5) with Intel Xeon Platinum 8380, Intel Xeon Gold 6348, or Intel Xeon Gold 6448H processors. We used R 4.2.3 [11] linked to Intel oneAPI Math Kernel Library (oneMKL) 2023.0.0.
