## Supplementary material for "Improving polygenic prediction from summary data by learning patterns of effect sharing across multiple phenotypes": S1 Table

Table 1: **Mean prediction $R^2$ across folds for the 16 blood cell traits in the full and sampled UK Biobank data.**

| **Phenotype** | **Full data** | | | **Sampled data** | | |
|---|---|---|---|---|---|---|
| | *LDpred2-auto* | *SBayesR* | *mr.mash-rss* | *LDpred2-auto* | *SBayesR* | *mr.mash-rss* |
| Red Blood Cell Counts (RBC#) | 0.1554 | 0.1577 | **0.1594** | 0.0883 | 0.0983 | **0.1043** |
| Haemoglobin Concentration (HGB) | 0.1162 | **0.1182** | 0.1164 | 0.0622 | **0.0711** | 0.0706 |
| Mean Corpuscular Volume (MCV) | 0.2273 | 0.2294 | **0.2325** | 0.1624 | 0.1727 | **0.1750** |
| Red Blood Cell Volume Distribution Width (RDW) | 0.1612 | 0.1635 | **0.1689** | 0.1028 | 0.1144 | **0.1230** |
| Mean Sphered Cell Volume (MSCV) | 0.1738 | 0.1765 | **0.1843** | 0.1129 | 0.1229 | **0.1322** |
| Reticulocyte Percentage (RET%) | 0.1366 | 0.1403 | **0.1452** | 0.0893 | 0.0991 | **0.1073** |
| High Light Scatter Reticulocytes Percentage (HLR%) | 0.1445 | 0.1482 | **0.1502** | 0.0959 | 0.1065 | **0.1115** |
| Platelet Count (PLT#) | 0.2385 | 0.2406 | **0.2428** | 0.1595 | 0.1702 | **0.1775** |
| Plateletcrit (PCT) | 0.1961 | 0.1982 | **0.1993** | 0.1213 | 0.1307 | **0.1346** |
| Platelet Distribution Width (PDW) | 0.2159 | 0.2182 | **0.2196** | 0.1524 | 0.1633 | **0.1661** |
| White Blood Cell Count (WBC#) | 0.1120 | 0.1135 | **0.1155** | 0.0564 | 0.0640 | **0.0667** |
| Lymphocyte Percentage (LYMPH%) | 0.0901 | 0.0917 | **0.0964** | 0.0422 | 0.0503 | **0.0561** |
| Monocyte Percentage (MONO%) | 0.1594 | 0.1624 | **0.1644** | 0.1081 | **0.1175** | 0.1159 |
| Neutrophil Percentage (NEUT%) | 0.0837 | 0.0858 | **0.0918** | 0.0402 | 0.0480 | **0.0546** |
| Eosinophil Percentage (EO%) | 0.1348 | **0.1365** | 0.1339 | 0.0714 | **0.0822** | 0.0802 |
| Basophil Percentage (BASO%) | 0.0213 | 0.0249 | **0.0283** | 0.0073 | 0.0143 | **0.0162** |

Highlighted in **bold** is the highest $R^2$ value across methods for each trait.
