## Supplementary material for "Improving polygenic prediction from summary data by learning patterns of effect sharing across multiple phenotypes": S2 Table

Table 2: **Mean $h_g^2$ across training sets for the 16 blood cell traits in the full UK Biobank data.**

| **Phenotype** | $\mathbf{h_g^2}$ |
|---|---|
| Red Blood Cell Counts (RBC#) | 0.23 |
| Haemoglobin Concentration (HGB) | 0.19 |
| Mean Corpuscular Volume (MCV) | 0.28 |
| Red Blood Cell Volume Distribution Width (RDW) | 0.22 |
| Mean Sphered Cell Volume (MSCV) | 0.23 |
| Reticulocyte Percentage (RET%) | 0.21 |
| High Light Scatter Reticulocytes Percentage (HLR%) | 0.22 |
| Platelet Count (PLT#) | 0.31 |
| Plateletcrit (PCT) | 0.26 |
| Platelet Distribution Width (PDW) | 0.24 |
| White Blood Cell Count (WBC#) | 0.20 |
| Lymphocyte Percentage (LYMPH%) | 0.16 |
| Monocyte Percentage (MONO%) | 0.20 |
| Neutrophil Percentage (NEUT%) | 0.16 |
| Eosinophil Percentage (EO%) | 0.20 |
| Basophil Percentage (BASO%) | 0.05 |
